# Doxorubicin-loaded human serum albumin nanoparticles overcome transporter-mediated drug resistance

**DOI:** 10.1101/655662

**Authors:** Hannah Onafuye, Sebastian Pieper, Dennis Mulac, Jindrich Cinatl, Mark N. Wass, Klaus Langer, Martin Michaelis

**Author notes:** equal contribution (Hannah Onafuye, Sebastian Pieper). to whom correspondence should be addressed: Klaus Langer, Institut für Pharmazeutische Technologie und Biopharmazie, WWU Münster, Corrensstrasse 48, 48149 Münster, Germany, / 83-39860, / 83-39308, Martin Michaelis, Industrial Biotechnology Centre and School of Biosciences, University of Kent, Canterbury CT2 7NJ, UK, / 82-7804, / 82-4034.

## Abstract

Resistance to systemic drug therapies is a major reason for the failure of anti-cancer therapies. Here, we tested doxorubicin-loaded human serum albumin (HSA) nanoparticles in the neuroblastoma cell line UKF-NB-3 and its ABCB1-expressing sublines adapted to vincristine (UKF-NB-3^r^VCR^1^) and doxorubicin (UKF-NB-3^r^DOX^20^). Doxorubicin-loaded nanoparticles displayed increased anti-cancer activity in UKF-NB-3^r^VCR^1^ and UKF-NB-3^r^DOX^20^ cells relative to doxorubicin solution, but not in UKF-NB-3 cells. UKF-NB-3^r^VCR^1^ cells were resensitised by nanoparticle-encapsulated doxorubicin to the level of UKF-NB-3 cells. UKF-NB-3^r^DOX^20^ cells displayed a more pronounced resistance phenotype than UKF-NB-3^r^VCR^1^ cells and were not re-sensitised by doxorubicin-loaded nanoparticles to the level of parental cells. ABCB1 inhibition using zosuquidar resulted in similar effects like nanoparticle incorporation, indicating that doxorubicin-loaded nanoparticles circumvent ABCB1-mediated drug efflux. The limited re-sensitisation of UKF-NB-3^r^DOX^20^ cells to doxorubicin by circumvention of ABCB1-mediated efflux is probably due to the presence of multiple doxorubicin resistance mechanisms. So far, ABCB1 inhibitors have failed in clinical trials, probably because systemic ABCB1 inhibition results in a modified body distribution of its many substrates including drugs, xenobiotics, and other molecules. HSA nanoparticles may provide an alternative, more specific way to overcome transporter-mediated resistance.

## Introduction

According to Globocan [1], there “were 14.1 million new cancer cases, 8.2 million cancer deaths and 32.6 million people living with cancer (within 5 years of diagnosis) in 2012 worldwide.” Despite substantial improvements over recent decades, the prognosis for many cancer patients remains unacceptably poor. The outlook is particularly grim for patients that are diagnosed with disseminated (metastatic) disease who cannot be successfully treated by local treatment (surgery, radiotherapy) and depend on systemic drug therapy, because the success of systemic therapies is typically limited by the occurrence of therapy resistance [2–4].

Drug efflux mediated by transporters including ATP-binding cassette (ABC) transporters has been shown to play a crucial role in cancer cell drug resistance [2,5]. ABCB1 (also known as P-glycoprotein or MDR1) seems to play a particularly important role in cancer cell drug resistance as a highly promiscuous transporter that mediates the cellular efflux of a wide range of structurally different substrates including many anti-cancer drugs. Different studies have reported that nano-sized drug carrier systems can bypass efflux-mediated drug resistance [6]. This includes various nanoparticle and liposome formulations of the ABCB1 substrate doxorubicin [7–12].

Here, we here investigated the effects of doxorubicin-loaded human serum albumin (HSA) nanoparticles in ABCB1-expressing neuroblastoma cells. HSA nanoparticles are easy to produce [13–17], and HSA is a well-tolerated material. It is the most abundant protein in human blood plasma and used in many pharmaceutical formulations, in particular as part of critical care treatment [18].

## Materials and methods

### Reagents and chemicals

HSA and glutaraldehyde were obtained from Sigma-Aldrich Chemie GmbH (Karlsruhe, Germany). Dulbecco’s Phosphate buffered saline (PBS) was purchased from Biochrom GmbH (Berlin, Germany). Doxorubicin was obtained from LGC Standards GmbH (Wesel, Germany). All chemicals were of analytical grade and used as received.

### Human serum albumin (HSA) nanoparticle preparation by desolvation

HSA nanoparticles were prepared by desolvation as previously described [13–17]. 100 μL of a 1% (w/v) aqueous doxorubicin solution were added to 500 μL of a 40 mg/mL (w/v) HSA solution and incubated for 2 h at room temperature under stirring (550 rpm, Cimaric i Multipoint Stirrer, ThermoFisher Scientific, Langenselbold, Germany). Then, 4 mL ethanol 96% were added at room temperature under stirring using a peristaltic pump (Ismatec ecoline, Ismatec, Wertheim-Mondfeld, Germany) at a flow rate of 1 mL/min. After the desolvation process, the resulting nanoparticles were stabilised/ cross-linked using different amounts of glutaraldehyde that corresponded to different percentages of the theoretic amount that is necessary for the quantitative crosslinking of the 60 primary amino groups present in the HSA molecules of the particle matrix. The addition of 4.7 μL 8% (w/v) aqueous glutaraldehyde solution resulted in a theoretical cross-linking of 40% of the HSA amino groups, the addition of 11.8 μL 8% (w/v) aqueous glutaraldehyde solution in 100% cross-linking, and the addition of 23.6 μL 8% (w/v) aqueous glutaraldehyde solution in 200% cross-linking. After glutaraldehyde addition, the suspension was stirred for 12 h at 550 rpm. The particles were purified by repeating three times centrifugation at 16,000 g for 12 min and resuspension in purified water. During particle purification the supernatants were collected, the drug content was measured by HPLC, and the loading efficiency of doxorubicin to the nanoparticles was calculated.

### Determination of particle size distribution

Average particle size and the polydispersity were measured by photon correlation spectroscopy (PCS) using a Malvern zetasizer nano (Malvern Instruments, Herrenberg, Germany). The resulting particle suspensions were diluted 1:100 with purified water and measured at a temperature of 22°C using a backscattering angel of 173°.

### Doxorubicin quantification via HPLC-UV

The amount of doxorubicin that had been incorporated into the nanoparticles was determined by HPLC-UV (HPLC 1200 series, Agilent Technologies GmbH, Böblingen, Germany) using a LiChroCART 250 × 4 mm LiChrospher 100 RP 18 column (Merck KGaA, Darmstadt, Germany). The mobile phase was a mixture of water and acetonitrile (70:30) containing 0.1% trifluoroacetic acid [16]. In order to obtain symmetric peaks a gradient was used. In the first 6 min the percentage of A was reduced from 70% to 50%. Subsequently within 2 min the amount of A was further decreased to 20% and then within another 2 min increased again to 70%. These conditions were held for a final 5 min resulting in a total runtime of 15 min. While using a flow rate of 0.8 mL/min, an elution time for doxorubicin of t = 7.5 min was achieved. The detection of doxorubicin was performed at a wavelength of 485 nm [19].

### Cell culture

The MYCN-amplified neuroblastoma cell line UKF-NB-3 was established from a stage 4 neuroblastoma patient [20]. UKF-NB-3 sub-lines adapted to growth in the presence of doxorubicin 20 ng/mL (UKF-NB-3^r^DOX^20^) [20] or vincristine 1 ng/mL (UKF-NB-3^r^VCR^1^) were established by continuous exposure to step-wise increasing drug concentrations as previously described [20,21] and derived from the Resistant Cancer Cell Line (RCCL) collection [22].

All cells were propagated in Iscove’s modified Dulbecco’s medium (IMDM) supplemented with 10% foetal calf serum, 100 IU/ml penicillin and 100 μg/ml streptomycin at 37°C. The drug-adapted sub-lines were continuously cultured in the presence of the indicated drug concentrations. Cells were routinely tested for mycoplasma contamination and authenticated by short tandem repeat profiling.

### Cell viability assay

Cell viability was determined by 3-(4,5-dimethylthiazol-2-yl)-2,5-diphenyltetrazolium bromide (MTT) assay modified after Mosman [23], as previously described [Michaelis et al., 24]. 2×10^4^ cells suspended in 100 μL cell culture medium were plated per well in 96-well plates and incubated in the presence of various drug concentrations for 120 h. Then, 25 μL of MTT solution (2 mg/mL (w/v) in PBS) were added per well, and the plates were incubated at 37°C for an additional 4 h. After this, the cells were lysed using 200 μL of a buffer containing 20% (w/v) sodium dodecylsulfate and 50% (v/v) N,N-dimethylformamide with the pH adjusted to 4.7 at 37°C for 4 h. Absorbance was determined at 570 nm for each well using a 96-well multiscanner. After subtracting of the background absorption, the results are expressed as percentage viability relative to control cultures which received no drug. Drug concentrations that inhibited cell viability by 50% (IC50) were determined using CalcuSyn (Biosoft, Cambridge, UK).

### Statistical testing

Results are expressed as mean ± S.D. of at least three experiments. Comparisons between two groups were performed using Student’s t-test. Three and more groups were compared by ANOVA followed by the Student-Newman-Keuls test. P-values lower than 0.05 were considered to be significant.

## Results

### Nanoparticle size, polydispersity and drug load

HSA nanoparticles were prepared by desolvation as previously described [13–17]. The nanoparticles were stabilised by the crosslinking of free amino groups present in albumin. Three different nanoparticle preparations were produced using glutaraldehyde at amounts that corresponded to a theoretical cross-linking of 40% (HSA 40% nanoparticles), 100% (HSA 100% nanoparticles), or 200% (HSA 200% nanoparticles) of the amino groups that are available in the HSA molecules. A non-stabilised (0% cross-linking) formulation was used as a control. The resulting particle sizes and polydispersity indices are shown in Table 1. HSA(0%) nanoparticles displayed a large particle size of almost 1 μm range and a high polydispersity of 0.5, confirming that no stable nanoparticles had formed (Table 1). The three HSA nanoparticle preparations stabilised by the different glutaraldehyde concentrations displayed similar diameters between 460 and 500 nm and polydispersity indices in the range of 0.153 and 0.213 indicating a narrow but not monodisperse size distribution (Table 1).

**Table 1.** Nanoparticle diameter, polydispersity, and drug load.

| Nanoparticles | Diameter<br>(nm) | Polydispersity | Drug load<br>($\mu\text{g}$ doxorubicin/ mg nanoparticle) |
| --- | --- | --- | --- |
| HSA(0%) | 848.7 | 0.500 | 370.9 |
| HSA(40%) | 485.8 | 0.189 | 151.9 |
| HSA(100%) | 496.4 | 0.213 | 190.5 |
| HSA(200%) | 463.4 | 0.153 | 164.8 |

While HSA(40%), HSA(100%), and HSA(200%) nanoparticles displayed similar drug loads between 152 and 191 μg doxorubicin/ mg nanoparticle, HSA(0%) nanoparticles had bound 371 μg doxorubicin/ mg HSA (Table 1). This probably reflected the higher accessibility of doxorubicin binding sites, which are known to be available on HSA [25], in HSA molecules in solution compared to the accessible binding sites available in HSA nanoparticles.

### Doxorubicin sensitivity of the used neuroblastoma cell lines

The parental neuroblastoma cell line UKF-NB-3 and its doxorubicin-(UKF-NB-3^r^DOX^20^) and vincristine-adapted (UKF-NB-3^r^VCR^1^) sub-lines substantially differed in their doxorubicin sensitivity (Figure 1). UKF-NB-3 displayed the lowest doxorubicin IC50 (3.8 ng/mL). UKF-NB-3^r^VCR^1^ was 4-fold more resistant to doxorubicin than UKF-NB-3 (doxorubicin IC50: 15.5 ng/mL). UKF-NB-3^r^DOX^20^ showed the highest doxorubicin IC50 (89.0 ng/mL) resulting in a 23-fold increase in doxorubicin resistance compared to UKF-NB-3 (Figure 1, Suppl. Table 1).

**Figure 1.**
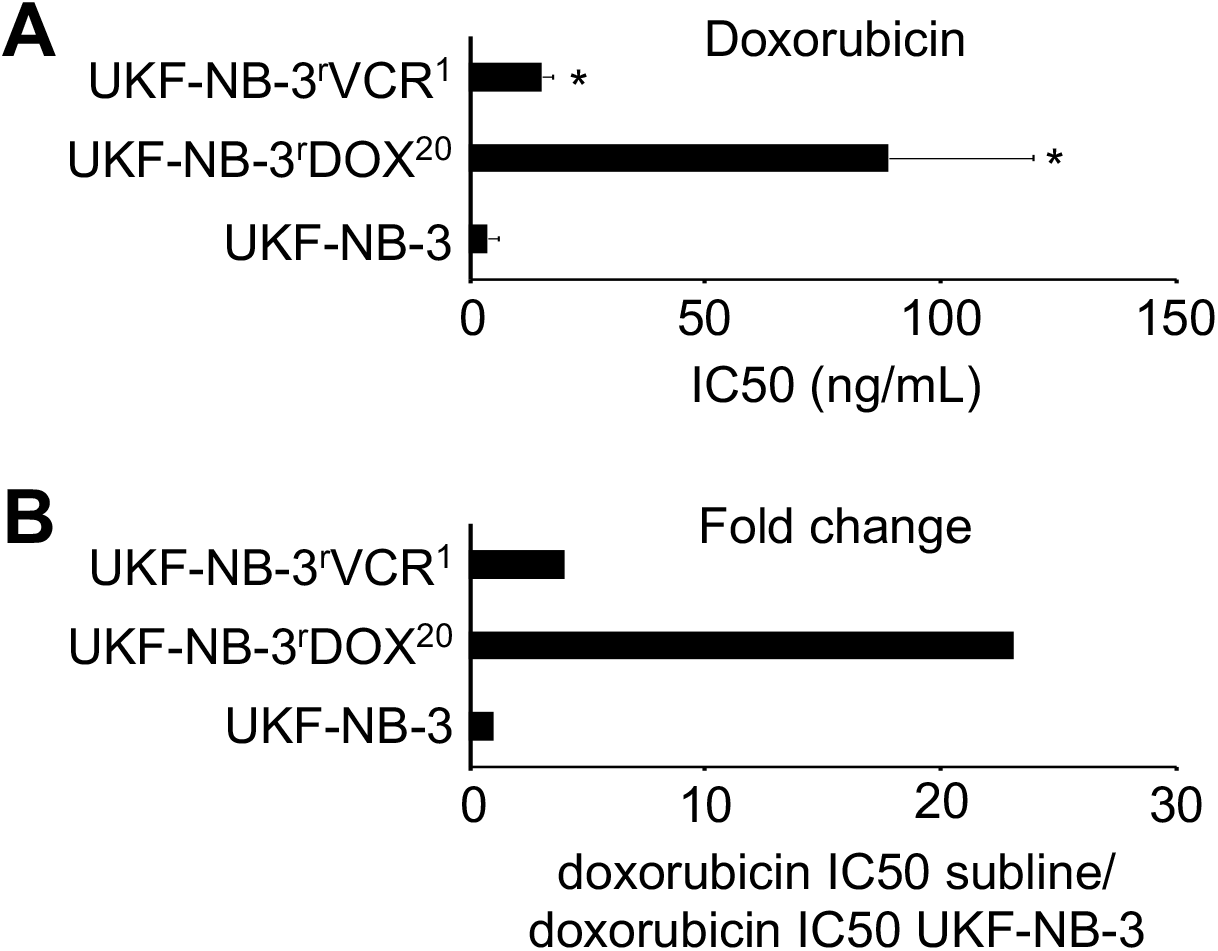
Doxorubicin sensitivity of UKF-NB-3, its doxorubicin-adapted sub-line UKF-NB-3^r^DOX^20^ and its vincristine-adapted sub-line UKF-NB-3^r^VCR^1^. A) Doxorubicin concentrations that reduce cell viability by 50% (IC50) as indicated by MTT assay after 120 h of incubation. B) Fold change in doxorubicin sensitivity (doxorubicin IC50 UKF-NB-3 sub-line/ doxorubicin IC50 UKF-NB-3). Numerical values are presented in Suppl. Table 1. * P < 0.05 relative to UKF-NB-3

### Effects of doxorubicin-loaded nanoparticles on neuroblastoma cells

The effects of doxorubicin applied in solution or incorporated into HSA(0%), HSA(40%), HSA(100%), or HSA(200%) nanoparticles on neuroblastoma cell viability are shown in Figure 2. The numerical values are presented in Suppl. Table 1. Empty control nanoparticles did not affect cell viability in the investigated concentrations.

**Figure 2.**
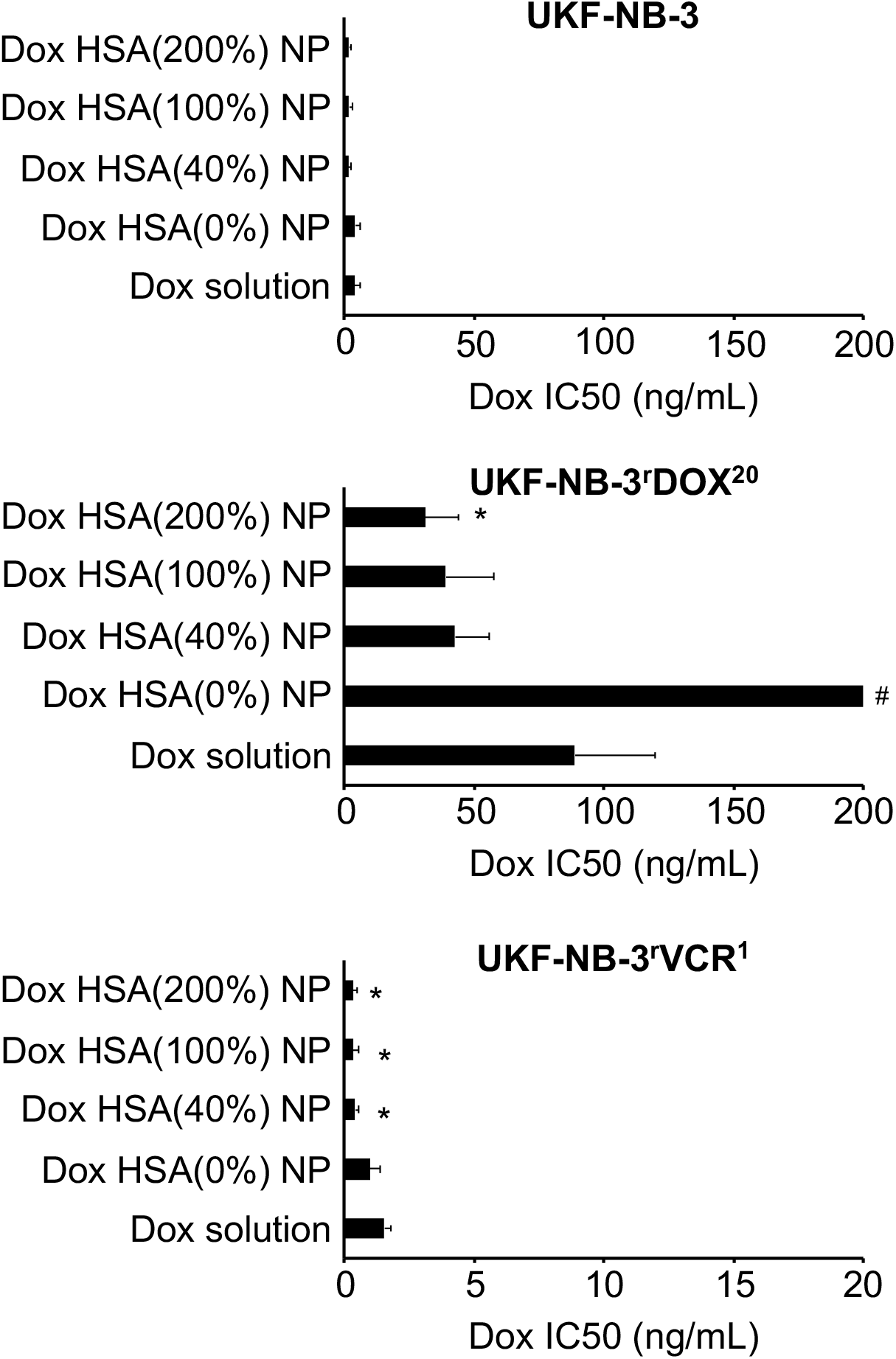
Effects of doxorubicin (Dox) applied as a solution or incorporated into human serum albumin (HSA) nanoparticles on neuroblastoma cell viability. The investigated nanoparticles differed in the amount of the cross-linker glutaraldehyde that was used for nanoparticle stabilisation. The amount of glutaraldehyde corresponded to 40% (Dox HSA(40%) NP), 100% (Dox HSA(100%) NP), or 200% (Dox HSA(200%) NP) theoretical cross-linking of the available amino groups present on HSA. Preparations prepared without glutaraldehyde served as a control (Dox HSA(0%) NP). Values are expressed as concentrations that reduce cell viability by 50% (IC50) as determined by MTT assay after 120 h of incubation. Numerical values are presented in Suppl. Table 1. Empty nanoparticles did not affect cell viability in the investigated concentrations. * P < 0.05 relative to doxorubicin solution; # IC50 > 200 ng/mL

In the neuroblastoma cell line UKF-NB-3, the nanoparticle preparations displayed similar activity as doxorubicin solution, with doxorubicin-loaded HSA(40%), HSA(100%), and HSA(200%) nanoparticles potentially showing a trend towards a slightly increased activity (Figure 2). However, the differences did not reach statistical significance. Similar results were obtained in the doxorubicin-adapted UKF-NB-3 sub-line UKF-NB-3^r^DOX^20^, although the difference between doxorubicin-loaded HSA(200%) nanoparticles and doxorubicin solution reached statistical significance (Figure 2). Notably, non-stabilised doxorubicin-bound HSA(0%) nanoparticles differed in their relative activity and did not reduce UKF-NB-3^r^DOX^20^ viability by 50% within the observed concentration range up to 200 ng/mL.

The vincristine-adapted UKF-NB-3 sub-line UKF-NB-3^r^VCR^1^ displayed decreased doxorubicin sensitivity. However, doxorubicin-loaded HSA(40%), HSA(100%), and HSA(200%) nanoparticles displayed a higher relative potency compared to doxorubicin solution in UKF-NB-3^r^VCR^1^ (Figure 2, Figure 3). The fold sensitisation doxorubicin IC50 doxorubicin solution/ doxorubicin IC50 nanoparticle-bound doxorubicin for HSA(40%), HSA(100%), and HSA(200%) nanoparticles (3.6 – 4.5-fold) was higher than for UKF-NB-3 (1.9 – 2.5-fold), and UKF-NB-3^r^DOX^20^ (2.1 – 2.9-fold). The differences between doxorubicin-loaded HSA(40%) nanoparticles, HSA(100%) nanoparticles, and HSA(200%) nanoparticles and doxorubicin solution reached statistical significance (P < 0.05) (Figure 2, Figure 3). Doxorubicin encapsulation into HSA(40%), HSA(100%), or HSA(200%) nanoparticles reduced the doxorubicin IC50 in UKF-NB-3^r^VCR^1^ cells to the levels of doxorubicin solution in parental UKF-NB-3 cells (Figure 2, Suppl. Table 1). In contrast, the doxorubicin IC50 of doxorubicin-loaded HSA nanoparticles remained clearly (8-11-fold) higher in UKF-NB-3^r^DOX^20^ cells than the doxorubicin IC50 of doxorubicin solution in parental UKF-NB-3 cells.

**Figure 3.**
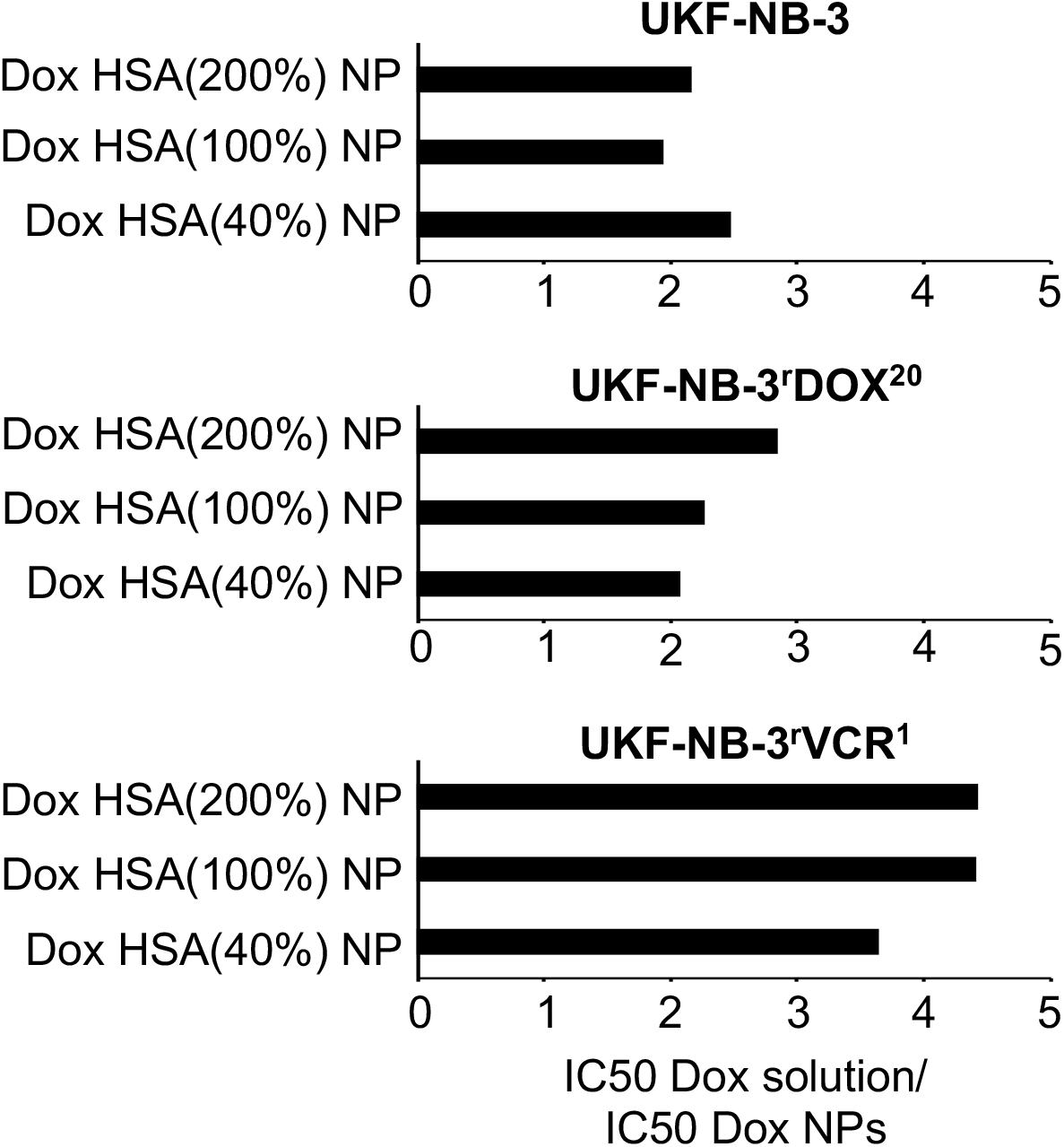
Fold sensitisation to doxorubicin by doxorubicin-bound nanoparticles (NP). Values are expressed as fold changes doxorubicin (Dox) IC50 of doxorubicin solution/ doxorubicin IC50 of doxorubicin-bound nanoparticles (NPs). Human serum albumin (HSA) nanoparticles were stabilised by glutaraldehyde concentrations corresponding to 40% (Dox HSA(40%) NP), 100% (Dox HSA(100%) NP), or 200% (Dox HSA(200%) NP) theoretical cross-linking of the available amino groups present on HSA.

### Effects of the ABCB1 inhibitor zosuquidar on the efficacy of nanoparticle-bound doxorubicin in UKF-NB-3^r^DOX^20^ cells

Doxorubicin is an ABCB1 substrate, and UKF-NB-3^r^DOX^20^ cells are characterised by high ABCB1 expression [20,26]. Vincristine is also an ABCB1 substrate, and vincristine-adapted cancer cell lines often display enhanced ABCB1 levels [20,26–29]. Accordingly, UKF-NB-3^r^VCR^1^ cells are sensitised by the ABCB1 inhibitor zosuquidar [2–6] to doxorubicin to the level of parental UKF-NB-3 cells (Suppl. Figure 1), which indicates that ABCB1 expression contributes to the resistance phenotype observed in UKF-NB-3^r^VCR^1^ cells.

Doxorubicin bound to nano-sized drug carrier systems has been shown to bypass ABCB1-mediated drug efflux [7–12]. In UKF-NB-3^r^VCR^1^ cells, both zosuquidar and doxorubicin encapsulation into HSA nanoparticles reduced the doxorubicin IC50 to the level of parental UKF-NB-3 cells (Figure 2, Suppl. Figure 1, Suppl. Table 1), which do not display detectable ABCB1 activity [20,27,29]. Hence, the increased activity of nanoparticle-bound doxorubicin that we observed in UKF-NB-3^r^VCR^10^ cells is likely to be attributed to the circumvention of ABCB1-mediated doxorubicin efflux.

In UKF-NB-3^r^DOX^20^ cells, however, the differences between doxorubicin solution and doxorubicin nanoparticles only reached statistical significance for doxorubicin-loaded HSA(200%) nanoparticles (Figure 2). Reasons for this may include that nanoparticle-incorporated doxorubicin do not completely avoid ABCB1-mediated efflux from UKF-NB-3^r^DOX^20^ cells and/ or that doxorubicin resistance is caused by multiple resistance mechanisms and that avoidance of ABCB1-mediated transport is not sufficient to re-sensitise UKF-NB-3^r^DOX^20^ cells to doxorubicin to the level of UKF-NB-3 cells.

To further study the role of ABCB1 as a doxorubicin resistance mechanism in UKF-NB-3^r^DOX^20^ cells, we performed additional experiments in which we combined the ABCB1 inhibitor zosuquidar and doxorubicin applied as a solution or nanoparticle preparations in UKF-NB-3^r^DOX^20^ and UKF-NB-3 cells. Zosuquidar (1 μM) did not affect the efficacy of doxorubicin solution or nanoparticle-bound doxorubicin in parental UKF-NB-3 cells (Figure 4), which do not display noticeable ABCB1 activity [20,27,29]. These experiments also confirmed that there is no significant difference in the anti-cancer activity between doxorubicin solution and doxorubicin nanoparticles in UKF-NB-3 cells, despite an apparent trend in the first set of experiments (Figure 2).

**Figure 4.**
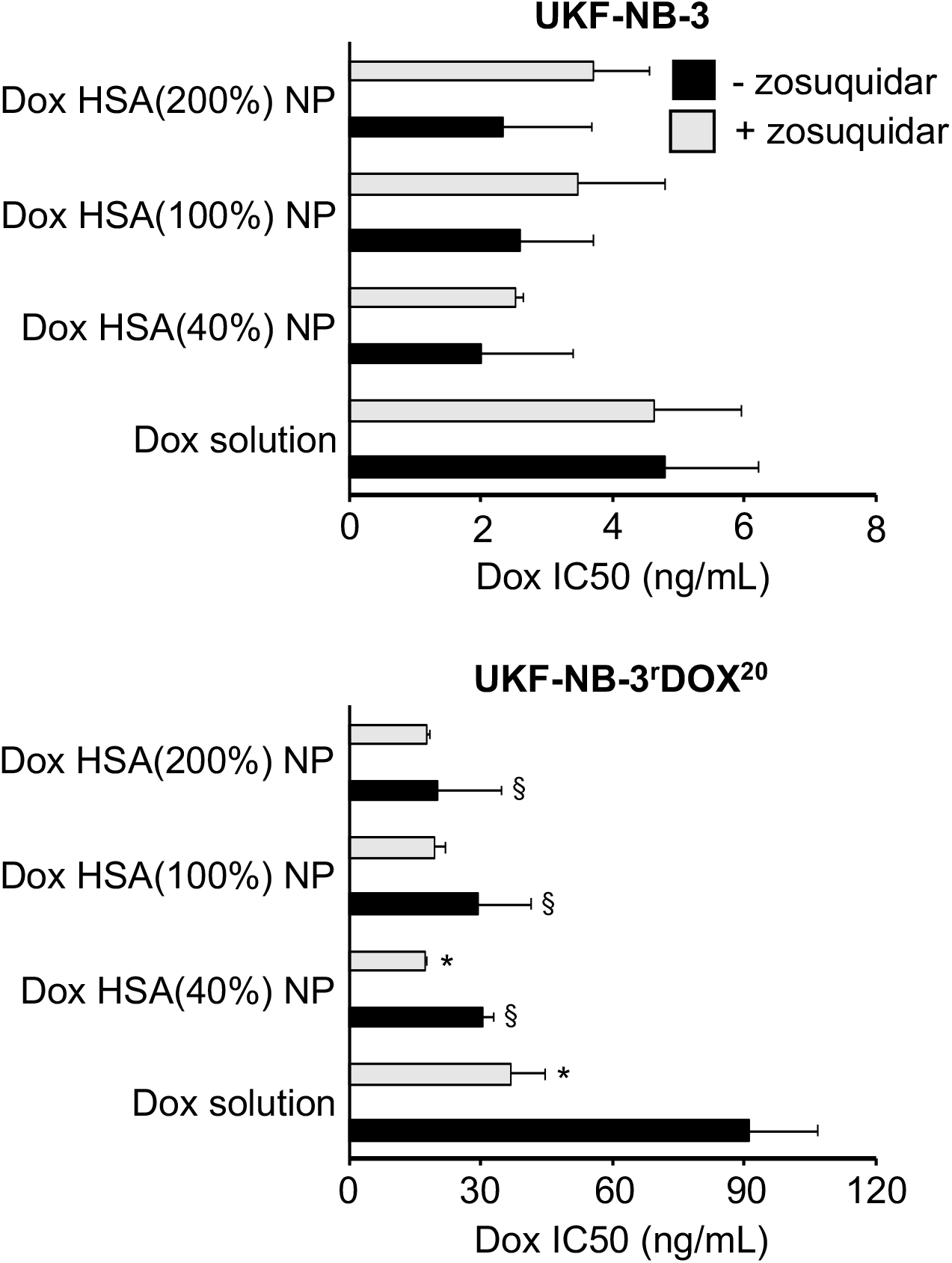
Doxorubicin (Dox) concentrations that reduce neuroblastoma cell viability by 50% (IC50) in the presence or absence of the ABCB1 inhibitor zosuquidar (1 μM) as determined by MTT assay after 120 h incubation. Doxorubicin was either applied as a solution or incorporated into human serum albumin (HSA) nanoparticles which had been stabilised by addition of glutaraldehyde concentrations corresponding to 40% (Dox HSA(40%) NP), 100% (Dox HSA(100%) NP), or 200% (Dox HSA(200%) NP) theoretical cross-linking of the available amino groups present on HSA. Zosuquidar (1 μM) did not affect cell viability on its own. Numerical data are presented in Suppl. Table 2. * P < 0.05 relative to the doxorubicin IC50 in the absence of zosuquidar; ^§^ P < 0.05 relative to doxorubicin solution

In UKF-NB-3^r^DOX^20^ cells, addition of zosuquidar resulted in an increased sensitivity to free doxorubicin (Figure 4). The doxorubicin IC50 decreased by 2.5-fold from 91 ng/mL in the absence of zosuquidar to 37 ng/mL in the presence of zosuquidar, but not to the level of UKF-NB-3 cells (4.6 ng/mL) (Suppl. Table 2). This confirmed that ABCB1 is one among multiple resistance mechanisms that contribute to the doxorubicin resistance phenotype observed in UKF-NB-3^r^DOX^20^.

In this set of experiments, doxorubicin-loaded nanoparticles displayed a significantly increased activity compared to doxorubicin solution in UKF-NB-3^r^DOX^20^ cells (Figure 4). This finding together with the non-significant trend observed in the first set of experiments (Figure 2) suggests that doxorubicin-loaded nanoparticles do indeed exert stronger effects against UKF-NB-3^r^DOX^20^ cells than doxorubicin solution. Zosuquidar only moderately increased the efficacy of doxorubicin nanoparticles further (1.1 – 1.8-fold) in UKF-NB-3^r^DOX^20^ cells (Figure 4, Suppl. Table 2). In particular, the anti-cancer effects of doxorubicin-loaded HSA(200%) nanoparticles, the most active nanoparticle preparation in UKF-NB-3^r^DOX^20^ cells, displayed a doxorubicin IC50 of 20 ng/mL, which was not further reduced by addition of zosuquidar (doxorubicin IC50: 18 ng/mL) (Figure 4, Suppl. Table 2). Hence, the increased anti-cancer activity of doxorubicin incorporated into HSA nanoparticles appears to be primarily caused by circumventing ABCB1-mediated doxorubicin efflux in UKF-NB-3^r^DOX^20^ cells.

## Discussion

The occurrence of drug resistance is the major reason for the failure of systemic anti-cancer therapies [2]. Here, we investigated the effects of doxorubicin-loaded HSA nanoparticles on the viability of the neuroblastoma cell line UKF-NB-3 and its sub-lines adapted to doxorubicin (UKF-NB-3^r^DOX^20^) and vincristine (UKF-NB-3^r^VCR^1^), which both display ABCB1 activity and resistance to doxorubicin. The HSA nanoparticles were prepared by desolvation and stabilised by glutaraldehyde, which crosslinks amino groups present in albumin molecules [13–17]. Glutaraldehyde was used at molar concentrations that corresponded to 40% (Dox HSA(40%) nanoparticles), 100% (Dox HSA(100%) nanoparticles), or 200% (Dox HSA(200%) nanoparticles) theoretical cross-linking of the amino groups available in the HSA molecules. The resulting nanoparticle preparations had similar sizes of about 200 nm and low polydispersity indices in the range of 0.2.

Doxorubicin-loaded nanoparticles displayed similar activity as doxorubicin solution in the parental UKF-NB-3 cell line, but exerted stronger effects than doxorubicin solution in the ABCB1-expressing UKF-NB-3 sub-lines. UKF-NB^r^VCR^1^ cells were similarly sensitive to doxorubicin-loaded nanoparticles as parental UKF-NB-3 cells to doxorubicin solution (and doxorubicin-loaded nanoparticles). This suggests that the doxorubicin resistance of UKF-NB-3^r^VCR^1^ cells exclusively depends on ABCB1 expression. In concordance, the ABCB1 inhibitor zosuquidar re-sensitised UKF-NB-3^r^VCR^1^ cells to the level of parental UKF-NB-3 cells.

UKF-NB-3^r^DOX^20^ cells displayed a more pronounced doxorubicin resistance phenotype than UKF-NB-3^r^VCR^1^ cells and were neither re-sensitised by nanoparticle-encapsulated doxorubicin nor by zosuquidar to the level of UKF-NB-3 cells. This suggests that UKF-NB-3^r^DOX^20^ cells have developed multiple doxorubicin resistance mechanisms. In contrast, adaptation of UKF-NB-3^r^VCR^1^ cells to vincristine, a tubulin-binding agent with an anti-cancer mechanism of action that is not related to that of the topoisomerase II inhibitor doxorubicin, did not result in the acquisition of changes that confer doxorubicin resistance beyond ABCB1 expression [2,20,30,31]. This indicates that the personalised use of nanoparticle-encapsulated transporter substrates will benefit from the use of biomarkers that indicate drug-specific resistance mechanisms in addition to transporter expression.

Furthermore, zosuquidar did not increase the efficacy of doxorubicin-loaded HSA(100%) and HSA(200%) nanoparticles and only modestly enhanced the efficacy of doxorubicin-loaded HSA(40%) nanoparticles. Together, these data confirm that administration of doxorubicin as HSA nanoparticles resulted in the circumvention of ABCB1-mediated drug efflux. The difference between HSA(40%) nanoparticles and the other two preparations may be explained by elevated drug release due to the lower degree of cross-linking.

Interestingly, high concentrations of the crosslinker glutaraldehyde did not affect the efficacy of the resulting doxorubicin-loaded nanoparticles although high glutaraldehyde concentrations might have been expected to affect drug release and/ or to bind covalently to doxorubicin via its amino group.

Notably, the results differ from a recent similar study in which nanoparticles prepared from poly(lactic-co-glycolic acid) (PLGA) or polylactic acid (PLA), two other biodegradable materials approved by the FDA and EMA for human use [32,33], did not bypass ABCB1-mediated drug efflux [34]. Differences in the mode of uptake and cellular distribution of nanoparticles from different materials may be responsible for these discrepancies. HSA nanoparticles may be internalised upon interaction with cellular albumin receptors [35,36]. Notably, nab-paclitaxel, an HSA nanoparticle-based preparation of paclitaxel (another ABCB1 substrate [26]), which is approved for the treatment of different forms of cancer [37], had previously been shown not to avoid ABCB1-mediated drug efflux [38]. However, nab-paclitaxel is not produced by the use of crosslinkers, and the interaction of paclitaxel with albumin may differ from that of doxorubicin. Hence, variations in drug binding and drug release kinetics may be responsible for this difference.

Despite the prominent role of ABCB1 as a drug resistance mechanism, attempts to exploit it as drug target have failed so far, despite the development of highly specific allosteric ABCB1 inhibitors (of which zosuquidar is one) [5,26]. A number of reasons seem to account for the clinical failure of ABCB1 inhibitors. ABCB1 is expressed at various physiological borders and involved in the control of the body distribution of its many endogenous and exogenous substrates. Systemic ABCB1 inhibition can therefore result in toxicity as consequence of a modified body distribution of anti-cancer drugs (and other drugs that are co-administered for other conditions than cancer), xenobiotics, and other molecules. In addition, cancer cells may be characterised by multiple resistance mechanisms (including the expression of multiple reporters) and targeting just one transporter may not be sufficient to overcome resistance (as supported by our current finding that UKF-NB-3^r^DOX^20^ cells cannot be fully re-sensitised to doxorubicin by zosuquidar) [2,5,26]. Hence, the use of drug carrier systems to bypass ABC transporter-mediated drug efflux is conceptually very attractive, because it can (in contrast to specific inhibitors of ABCB1 or other transporters) overcome resistance mediated by multiple transporters and does not result in the systemic inhibition of ABC transporter function at physiological barriers.

In conclusion, doxorubicin-loaded HSA nanoparticles produced by desolvation and crosslinking using glutaraldehyde overcome (in contrast to other nanoparticle systems) transporter-mediated drug resistance.

## Acknowledgements

This work was supported by the Kent Cancer Trust, the Hilfe für krebskranke Kinder Frankfurt e.V., and the Frankfurter Stiftung für krebskranke Kinder.

---

**Suppl. Figure 1.**
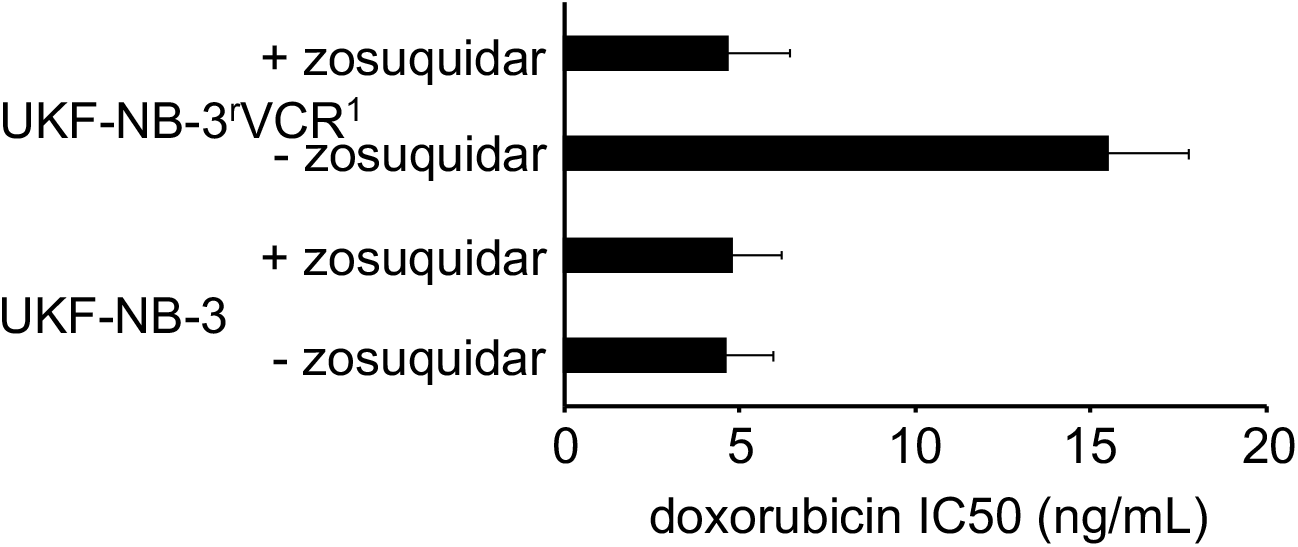
Doxorubicin concentrations that reduce neuroblastoma cell viability by 50% (IC50) in the absence or presence of the ABCB1 inhibitor zosuquidar (1 μM).

**Suppl. Table 1.**
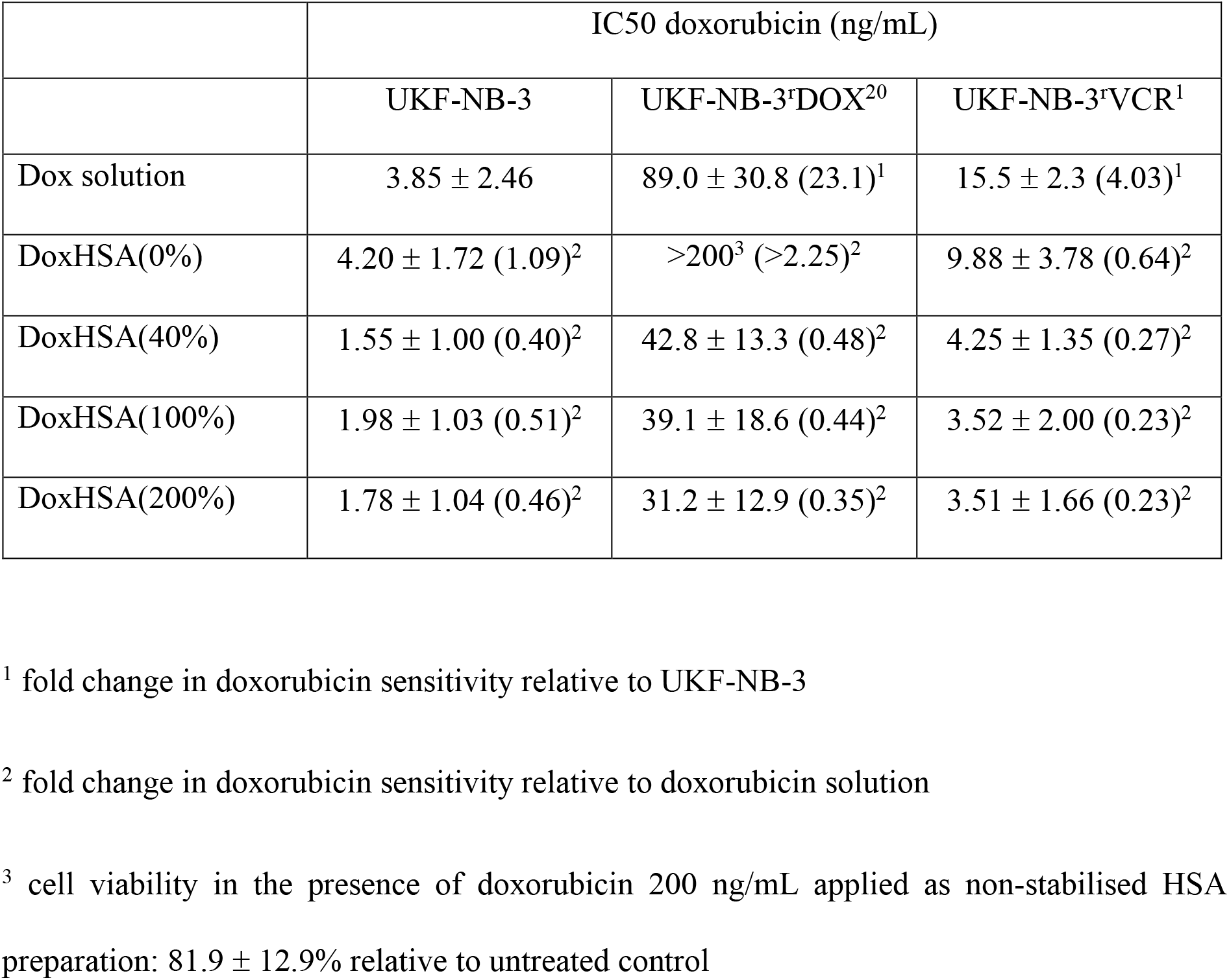
Effects of doxorubicin (Dox) applied as solution or incorporated into human serum albumin (HSA) nanoparticles on neuroblastoma cell viability. The investigated nanoparticles differed in the amount of the crosslinker glutaraldehyde that was used for nanoparticle stabilisation. The glutaraldehyde amount corresponded to 40% (Dox HSA(40%) NP), 100% (Dox HSA(100%) NP), or 200% (Dox HDA(200%) NP) of the theoretical amount of available amino groups present on HSA. Preparations prepared without glutaraldehyde served as control (Dox HSA(0%) NP). Values are expressed as concentrations that reduce cell viability by 50% (IC50) as determined by MTT assay after 120h of incubation.

**Suppl. Table 2.**
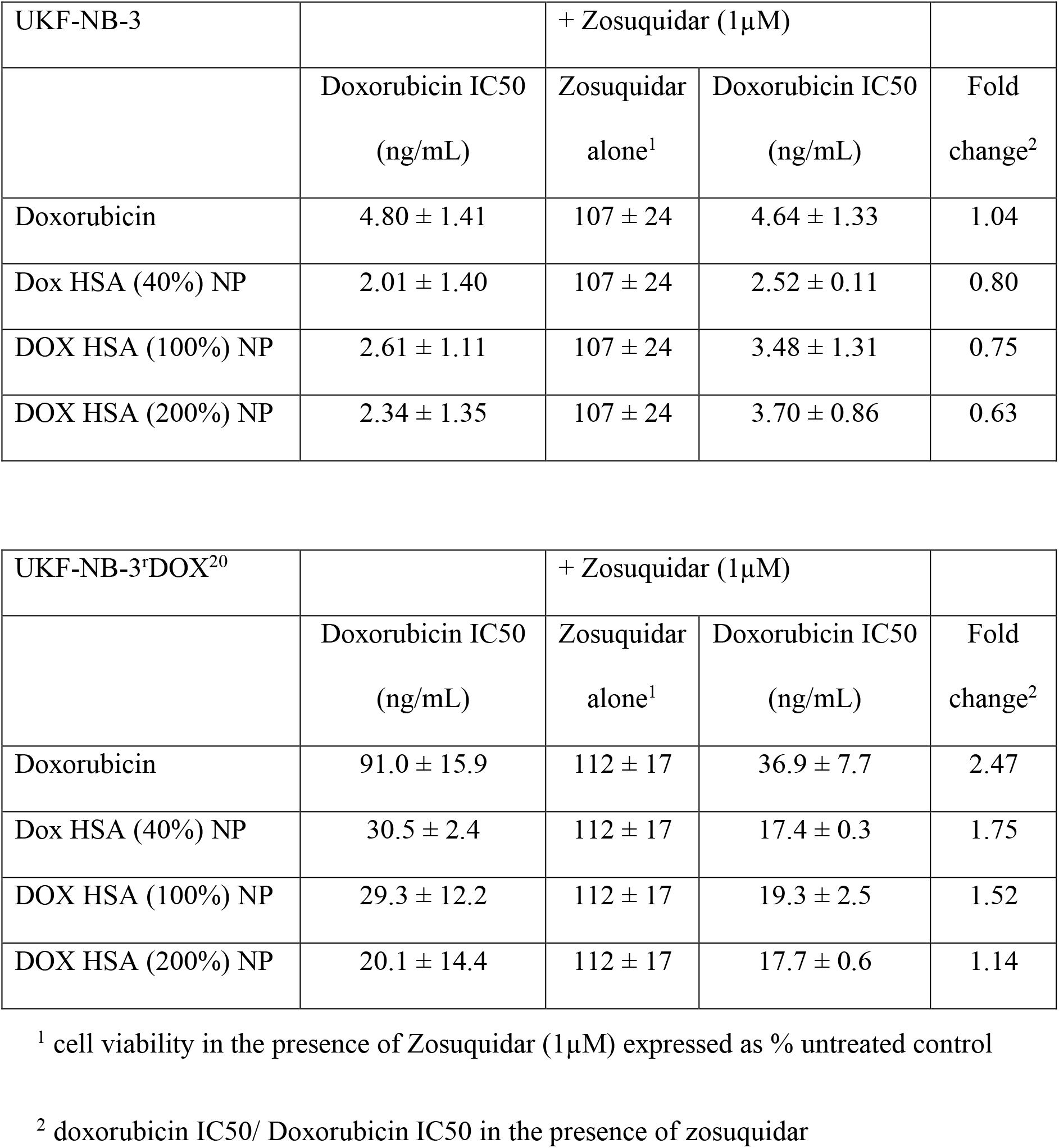
Effects of doxorubicin (Dox) applied as solution or incorporated into human serum albumin (HSA) nanoparticles on neuroblastoma cell viability in the absence or presence of zosuquidar (1μM). The investigated nanoparticles differed in the amount of the crosslinker glutaraldehyde that was used for nanoparticle stabilisation. The glutaraldehyde amount corresponded to 40% (Dox HSA(40%) NP), 100% (Dox HSA(100%) NP), or 200% (Dox HDA(200%) NP) of the theoretical amount of available amino groups present on HSA.. Values are expressed as concentrations that reduce cell viability by 50% (IC50) as determined by MTT assay after 120h of incubation.

